# Inhalation: A means to explore and optimize nintedanib’s pharmacokinetic/pharmacodynamic relationship

**DOI:** 10.1101/2020.03.27.012401

**Authors:** Gali Epstein-Shochet, Stephen Pham, Steven Beck, Safaa Naiel, Olivia Mekhael, Spencer Revill, Aaron Hayat, Megan Vierhout, Becky Bardestein-Wald, David Shitrit, Kjetil Ask, A. Bruce Montgomery, Martin R. Kolb, Mark W. Surber

**Author notes:** Corresponding author: Mark W. Surber, Avalyn Pharma, Inc., 2251 San Diego Ave, B-111, San Diego, CA 92110, United States.

## Abstract

Oral nintedanib is marketed for the treatment of idiopathic pulmonary fibrosis (IPF). While effective slowing fibrosis progression, as an oral medicine nintedanib is limited. To reduce side effects and maximize efficacy, nintedanib was reformulated as a solution for nebulization and inhaled administration. To predict effectiveness treating IPF, the nintedanib pharmacokinetic/pharmacodynamic relationship was dissected. Pharmacokinetic analysis indicated oral-delivered nintedanib plasma exposure and lung tissue partitioning were not dose-proportional and resulting lung levels were substantially higher than blood. Although initial-oral absorbed nintedanib efficiently partitioned into the lung, only a quickly eliminated fraction appeared available to epithelial lining fluid (ELF). Because IPF disease appears to initiate and progress near the epithelial surface, this observation suggests short duration nintedanib exposure (oral portion efficiently partitioned to ELF) is sufficient for IPF efficacy. To test this hypothesis, exposure duration required for nintedanib activity was explored. *In vitro*, IPF-cellular matrix (IPF-CM) increased primary normal human fibroblast (nHLF) aggregate size and reduced nHLF cell count. IPF-CM also increased nHLF ACTA2 and COL1A expression. Whether short duration (inhalation mimic) or continuous exposure (oral mimic), nintedanib (1-100 nM) reversed these effects. *In vivo*, intubated silica produced a strong pulmonary fibrotic response. Once-daily (QD) 0.021, 0.21 and 2.1 mg/kg intranasal (IN; short duration inhaled exposure) and twice daily (BID) 30 mg/kg oral (PO; long duration oral exposure) showed that at equivalent-delivered lung concentrations, QD short duration inhaled nintedanib exposure (0.21 mg/kg IN vs. 30 mg/kg PO) exhibited equivalent-to-superior activity as BID oral (reduced silica-induced elastance, alpha-smooth muscle actin, interleukin-1 beta (IL-1β) and soluble collagen, and lung macrophage and neutrophils). An increased lung dose (2.1 mg/kg IN vs. 30 mg/kg PO) exhibited increased effect by further reducing silica-induced elastance, IL-1β and soluble collagen. Neither oral nor inhaled nintedanib reduced silica-induced parenchymal collagen. Both QD inhaled and BID oral nintedanib reduced silica-induced inflammatory index with oral achieving significance. In summary, nintedanib pulmonary anti-fibrotic activity can be achieved using small, infrequent inhaled doses to deliver oral equivalent-to-superior therapeutic effect.

## 1. INTRODUCTION

Idiopathic pulmonary fibrosis (IPF) is a disease occurring primarily in older adults, characterized by chronic loss of pulmonary capacity and poor prognosis ^[1, 2]^. IPF is a specific form of chronic fibrosing interstitial pneumonia limited to the lung and associated with the pathological pattern of usual interstitial pneumonia. IPF follows a variable clinical course in which periods of relative stability are mixed with episodes of accelerated decline, possibly resulting in respiratory failure and death.

Much of IPF pathogenesis remains to be understood. However, it is believed that interactions between airway and alveolar epithelial cells with underlying fibroblasts are important for disease initiation and progression ^[3]^. Evidence includes recurrent and/or non-resolving epithelial injury stimulating fibroblast accumulation and differentiation into myofibroblasts; accumulated myofibroblasts centralize into fibroblastic foci (regions of excess collagen production); and the close apposition of airway and alveolar epithelial cells with fibroblastic foci. Together, this relationship supports the airway and alveolar surface as the targeted location for therapeutic IPF drug delivery.

Pro-fibrotic tyrosine kinases such as platelet-derived growth factor receptor (PDGFR) and fibroblast growth factor receptor (FGFR) have been implicated in the pathogenesis of IPF ^[4-6]^. As key regulators of fibroblast proliferation and migration in the lung, these and other tyrosine kinase receptors play an important role in IPF disease initiation and progression. Nintedanib is a small-molecule tyrosine kinase inhibitor that competitively blocks the adenosine triphosphate binding pocket of both receptor tyrosine kinases and non-receptor tyrosine kinases, such as IPF-implicated PDGFR, FGFR ^[7]^ and Src (proto-oncogene tyrosine-protein kinase src) kinase families ^[8, 9]^. Blocking these kinase activities impacts downstream signalling pathways crucial for normal and disease-associated cellular activity ^[6, 10]^. Based upon one phase 2 ^[11]^ and two phase 3 registration clinical trials ^[12]^, each demonstrating a significant reduction in the annual rate of forced vital capacity (FVC) decline (consistent with slowed disease progression), oral nintedanib was approved for the treatment of IPF (marketed as Ofev^®^).

In the phase 2 study, patients received oral nintedanib as either 50 mg once a day (QD), 50 mg twice a day (BID), 100 mg BID, 150 mg BID or placebo for 52 weeks. Only the 150 mg BID dose showed benefit, with a significant 68.4% reduction in rate of annual FVC decline. The most frequent adverse events in this group were diarrhea (55.3%) and increased aminotransferase levels at least three times the upper limit of the normal range (7.1%). In the phase 3 studies, patients received either 150 mg BID oral nintedanib or placebo for 52 weeks. The two studies showed a significant 52.2% and 45.2% reduction in rate of annual FVC decline. The most frequent adverse events were diarrhea (61.5% and 63.2%) and increased aminotransferase levels at least three times the upper limit of the normal range (4.9% and 5.2%). Of treated patients, 26.5% and 29.2% were dose-reduced from 150 mg BID to 100 mg BID, and 25.6% and 21.9% of patients had their treatment interrupted for a mean duration of 3.9 and 3.2 weeks, respectively. Less than 5% of patients discontinued study medication. In 2017, the Ofev label was updated to include new risks of liver-induced injury, hyperbilirubinemia, jaundice, and drug-induced liver injury (Ofev label).

As shown in the phase 2 study (and studied in phase 3), only the highest oral nintedanib dose showed effect. Unfortunately, substantial side effects prevented further oral dose escalation for possible additional efficacy. Complicating matters, first-pass metabolism, and safety driven dose reduction and stoppage protocols further reduce lung dose and interrupt required maintenance therapy (Ofev Label). To overcome these shortcomings and maximize efficacy, nintedanib was reformulated as a solution for nebulization and inhaled, direct-lung administration. To compare the effectiveness of inhaled nintedanib to oral, a rat bleomycin model delivering similar inhaled and oral lung exposures resulted in equivalent anti-fibrotic efficacy ^[13]^. In the same model, inhaled dose levels delivering higher nintedanib lung exposures resulted in superior efficacy and improved animal weight gain to levels equivalent to sham animals; a result far superior to stunted weight gain associated with oral administration.

Because the inhaled approach holds promise to circumvent oral adverse events and improve efficacy, we explored the oral pharmacokinetic/pharmacodynamic relationship to better predict inhaled nintedanib effectiveness in treating IPF. In this report, we provide several models comparing the effect of oral and inhaled pharmacokinetics and discuss the impact of nintedanib exposure duration on nintedanib *in vitro* and *in vivo* activity.

## 2. MATERIALS AND METHODS

### 2.1 Materials

Nintedanib base was purchased from Tecoland (Irvine, California). Nintedanib esylate was provided by Fermion Oy (Espoo, Finland). Alternative salt forms (nintedanib HCl and nintedanib HBr) were provided by Avalyn Pharma. Dose solution compositions may be found in the Supplement Table S-1 ^[14]^. Briefly, for cell culture, nintedanib base was dissolved in DMSO. Oral (PO) dose solutions were prepared in water with 1% methylcellulose. Inhaled dose solutions for intratracheal aerosol (IT) administration were prepared in water with 2% propylene glycol and delivered using a Penn Century Aerosolizer® device (Wyndmoor, Pennsylvania). Inhaled dose solutions for intranasal (IN) administration were prepared in water with 1.5% propylene glycol and 0.4% sodium chloride. Silica was obtained from U.S. Silica (Katy, Texas). Isoflurane anesthesia was obtained from MTC Pharmaceuticals (Cambridge, Ontario).

### 2.2 Pharmacokinetic analysis of inhaled and oral nintedanib

Mice were administered nintedanib esylate by PO or IT delivery. PO and IT pharmacokinetic studies were performed at SMC Laboratories, Inc. (Tokyo, Japan). Six-week-old female C57BL/6 mice (18-20 grams) were obtained from SLC, Inc. (Japan). Animals were maintained under controlled conditions of temperature (23 ± 2°C), humidity (45 ± 10%) and lighting (12-hour artificial light and dark cycles) with air exchange. Animals were housed in TPX cages (CLEA Japan) with a maximum of 6 mice per cage. Sterilized Paper-Clean (Japan SLC) was used for bedding. Sterilized normal diet (CE-2; CLEA, Japan) and purified water were provided *ad libitum*. After acclimation, animals were randomized into groups of 24 mice.

For PO administration, nintedanib (as nintedanib esylate) was dosed in 10 mL/kg at 10 and 100 mg/kg. For IT administration, nintedanib (as nintedanib esylate, nintedanib HCl and nintedanib HBr) was dosed at 1, 2.5 and 10 mg/kg. Because IT pharmacokinetic parameters between the three nintedanib salt forms were determined equivalent, results were pooled (n of 9 animals per time point) and described herein as nintedanib. Inclusion of permeant anion (e.g., chloride) is usually desired for aerosol upper airway tolerability. However, because nintedanib exhibits poor long-term stability in chloride-containing solutions and IT administration by-passes the upper airway, permeant anion-containing salts were excluded from IT dosing solutions. To maintain osmotic balance, propylene glycol was included. IT nintedanib delivered 50 μL formulation directly to the lung by nebulizing catheter as described previously ^[15]^. Plasma and lung tissue samples were taken 2, 5, 10, 20, 40 min and 1, 2, and 4 hours post dose. At designated collection times, blood (100 μL) was sampled to prepare K3EDTA plasma and lungs were collected. Briefly (all times post-dose; n of 9 animals per IT dose level per time point, n of 3 animals per oral dose level per time point): Group 1: blood at 2 min and 40 min, lungs at 40 min; Group 2: blood at 5 min and 60 min, lungs at 60 min; Group 3: blood at 10 min and 120 min, lungs at 120 min; Group 4: blood at 20 min and 240 min, lung at 240 min; Group 5: lungs only 2 min; Group 6: lungs only at 5 min; Group 7: lungs only at 10 min; Group 8: lungs only at 20 min. Within each group, initial blood samples were taken from the facial vein. Final blood samples were collected, and exsanguination was performed through the abdominal aorta under sodium pentobarbital anesthesia (Kyoritsu Seiyaku, Japan). Bronchoalveolar lavage (BAL; n of 3 PO animals and n of 9 IT animals per time point) was performed 120 and 240-min post-PO and 60 and 180-min post-IT administration.

Previous multi-dose mouse and rat bleomycin efficacy studies indicated IT administration was not well-tolerated (IT requires deep anesthetic limiting animal appetite and growth; data not shown). To optimize animal health over the study period, we elected to perform the silica-induced pulmonary fibrosis study using IN administration, requiring only light anesthetic and further avoiding possible tracheal injury associated with IT administration. As confirmation, a tolerability study was first performed in silica-challenged mice (see Supplement ^[14]^). For IN dose selection, a pharmacokinetic bridging study comparing IT administration was also performed (see Supplement).

Plasma, lung and BAL fluid (BALF) bioanalytical sample analysis was performed by MicroConstants, Inc, (San Diego, USA) as described in the Supplement ^[14]^. Pharmacokinetic parameters were determined using the linear trapezoidal method.

### 2.3 Primary human lung fibroblast (HLF) culture

Primary human lung fibroblasts were isolated from at least six IPF (histologically confirmed) tissues and 10 control samples (histologically normal lung taken during lobectomy distant from the resected tumor), as described ^[16]^. Tissue was obtained at the time of biopsy. Following extraction, fibroblasts were cultured in Dulbecco’s Modified Eagle’s Medium (DMEM) supplemented with 20% FCS, L-glutamine (2 mM), and antibiotics (Biological Industries, Israel) and maintained in 5% CO2 at 37°C. Harvested human lung fibroblasts from both IPF patients (IPF-HLFs) and control donors (nHLFs) had typical spindle morphology and were alpha smooth muscle actin (αSMA) positive.

### 2.4 IPF-conditioned cellular matrix (IPF-CM) model

Experiments were performed as previously described ^[17]^. Briefly, IPF-HLFs or nHLFs were cultured in 24-well plates previously coated with 10 mg/mL growth factor-reduced (GFR) Matrigel (BD Biosciences, San Jose, California). Following 48 h, cells were removed by NH4OH. nHLFs (15×10e4) were either exposed to nintedanib (1-100 nM) for 60 min, washed and then added for further culture with the same nintedanib concentration or exposed to nintedanib (1-100 nM) for 60 min, washed and then added for further culture without nintedanib. Following an additional 24 h incubation, cultures were examined using Olympus IX71 microscope. Aggregate size and number were measured by ImageJ: http://rsbweb.nih.gov/ij/. To accommodate both increased aggregate size and the decreased total aggregate number, average aggregate size was calculated.

### 2.5 RNA extraction and qPCR

After culturing, cells were harvested for total RNA extraction using the RNeasy kit (QAIGEN, USA). RNA was converted to cDNA using GeneAmp (Applied Biosystems, USA). Reactions were performed using Power SYBR Green (Applied Biosystems, UK). Primer sequences (purchased from Hylabs, Israel) are listed in Table 1.

**Table 1.**
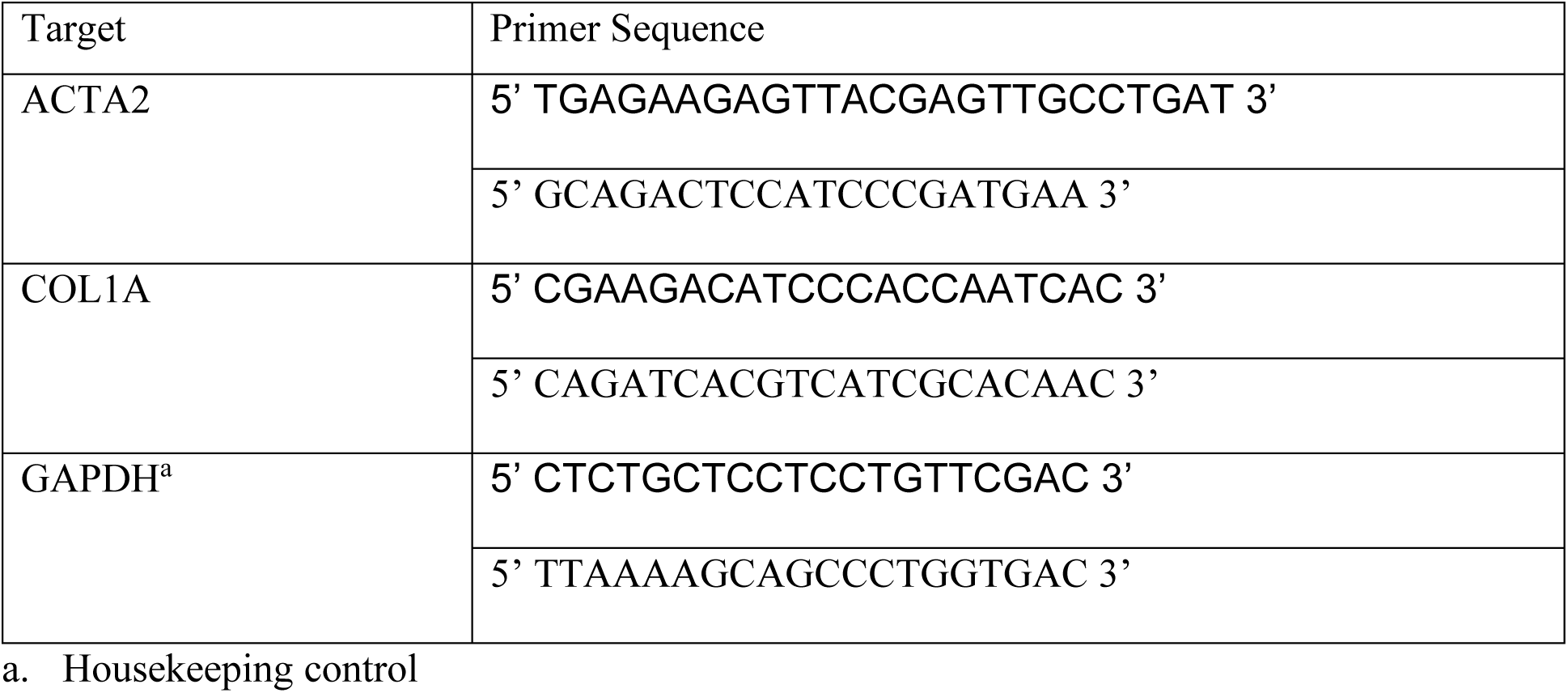
qPCR Primer Sequences.

### 2.6 Therapeutic silica-induced pulmonary fibrosis model

To explore the nintedanib pharmacokinetic relationship with efficacy, a modified version of the mouse silica model was used ^[18]^. Briefly, while under light isoflurane anesthesia (MTC Pharmaceuticals, Cambridge, ON, Canada), experimental pulmonary fibrosis was induced in 20-22 g female C57BL/6 mice (Charles River Laboratories) by a single silica IT intubation (2.5 mg/kg) or PBS vehicle control. On day 10 through day 29, groups received either PO or IN vehicle or nintedanib treatment. IN animals were lightly anesthetized with isoflurane and dosed once daily (QD) with 0.021, 0.21 or 2.1 mg/kg nintedanib (35 μL per dose). PO animals were not anesthetized and dosed twice daily (BID). As previously shown effective ^[10]^, PO-treated animals were dosed at 30 mg/kg BID (10 mL/kg).

Five animals were enrolled into sham groups, ten into each PO dose group and thirteen into each IN dose group. Compared to oral lung levels, IN nintedanib dose levels were selected to deliver inferior, equivalent and superior lung Cmax and AUC (Supplement Table S-2 and Figure S-1 ^[14]^). All animals were euthanized on day 30. Each mouse was weighed at the beginning and throughout the study; weight fluctuations were represented as a percent change from starting weight. Flexivent was performed to determine elastance (lung function). Left lungs were extracted, stained for parenchymal collagen and αSMA, while right lung homogenates were assessed for soluble collagen and interleukin-1β (IL-1β). Method details may be found in the Supplement ^[14]^.

### 2.7 Statistical analysis

Statistical analysis was performed using GraphPad Prism 8.0d (GraphPad Software, Inc). The Rout’s method was performed to identify outliers. When two groups were compared, a normality test was performed before the appropriate T-test. For more than two groups, data were analyzed using a one-way ANOVA with a Dunnett multiple comparisons test (post hoc). A p-value less than 0.05 was considered significant. Percent benefit analysis was performed by first subtracting sham animal results from vehicle and treatment results, then determining the percent difference between remaining treatment and vehicle values.

### 2.8 Ethical approval

All IT and PO pharmacokinetic study animals were housed and cared for in accordance with the Japanese Pharmacological Society Guidelines for Animal Use. The IN pharmacokinetic, as well as IN and PO efficacy studies were sanctioned and approved by the Animal Research Ethics Board of McMaster University and carried out in accordance to the guidelines of the Canadian Council on Animal Care. Animals were monitored for weight loss and other signs of physical distress throughout these experiments. The primary human lung sample study was approved by the institutional Ethics Committee of Meir Medical Center (study number 0016-16-MMC). Informed consent was obtained from all patients.

## 3. RESULTS

### 3.1 Nintedanib pharmacokinetics and lung-tissue distribution

Mice were administered nintedanib by PO or IT delivery. Plasma and lungs were collected at various time points and nintedanib was extracted and quantified as ng/mL plasma and µg/gram homogenized lung tissue. Results are shown in Table 2.

**Table 2.** Nintedanib pharmacokinetics following oral and intratracheal administration.

|  |  |  | Oral (SEM) |  | Inhaled (SEM) |  |  |  |
| --- | --- | --- | --- | --- | --- | --- | --- | --- |
|  |  |  | Gavage |  | IT<sup>a</sup> |  |  | IN<sup>b</sup> |
| Dose mg/kg |  |  | 10 | 100 | 1 | 2.5 | 10 | 3 |
| Plasma | AUC<sub>0-last</sub> | mg·hr/L | 0.1<br>(0.03) | 5.5<br>(1.16) | 0.2<br>(0.05) | 0.2<br>(0.05) | 0.7<br>(0.13) | 0.3<br>(0.09) |
|  | C<sub>max</sub> | ng/mL | 12<br>(3.0) | 962<br>(202.0) | 320<br>(80.0) | 1,270<br>(292.1) | 4,860<br>(874.8) | 2,140<br>(611.4) |
|  | T<sub>max</sub> | min | 246<br>(2.3) | 230<br>(6.0) | 0 | 0 | 0 | 0 |
|  | T<sub>1/2α</sub> | min | n/a | n/a | 1.9<br>(0.28) | 2.2<br>(0.46) | 0.7<br>(0.13) | 5.2<br>(1.55) |
|  | T<sub>1/2term</sub> | min | 65<br>(16.25) | 60<br>(11.4) | 121<br>(24.2) | 126<br>(21.4) | 153<br>(19.9) | 124<br>(8.1) |
| Lung<sup>c</sup> | AUC<sub>0-last</sub> | mg·hr/kg | 2.3<br>(1.2) | 77.6<br>(9.2) | 28.3<br>(3.9) | 50.1<br>(5.9) | 148.5<br>(4.2) | 56.9<br>(8.5) |
|  | C<sub>max</sub> | μg/g | 0.4<br>(0.2) | 13<br>(1.5) | 48<br>(6.6) | 167<br>(19.5) | 441<br>(12.3) | 42<br>(6.3) |
|  | T<sub>max</sub> | min | 294<br>(10.4) | 318<br>(10.0) | 0 | 0 | 0 | 0 |
|  | T<sub>1/2α</sub> | min | n/a | n/a | 1.6<br>(0.2) | 1.4<br>(0.2) | 1.4<br>(0.0) | 14.6<br>(2.3) |
|  | T<sub>1/2term</sub> | min | 108<br>(35.6) | 131<br>(14.5) | 101<br>(13.3) | 102<br>(9.2) | 125<br>(13.0) | 202<br>(24.9) |

a. IT: Intratracheal; b. IN: Intranasal; C. Lung: Whole lung homogenate

Compared to 10 mg/kg PO, results indicate 100 mg/kg PO-administered nintedanib delivered an 80-fold greater plasma Cmax and 55-fold greater plasma AUC. Lung tissue homogenate results indicate 100 mg/kg PO delivered 33-fold greater lung Cmax and 34-fold greater lung AUC.

Compared to PO, 1.0, 2.5 and 10 mg/kg IT administered nintedanib delivered 2, 2 and 7-fold greater plasma AUC, respectively than 10 mg/kg PO, and 0.04, 0.04 and 0.1-fold compared to 100 mg/kg PO. Delivered plasma Cmax from the same IT dose levels were 27, 105 and 405-fold greater than 10 mg/kg PO, and 0.3, 1.3 and 5-fold compared to 100 mg/kg PO. Lung tissue homogenate results indicated 1.0, 2.5 and 10 mg/kg IT administered nintedanib delivered 12, 22 and 65-fold greater lung AUC, respectively than 10 mg/kg PO, and 0.4, 0.7 and 2-fold compared to 100 mg/kg PO. Delivered lung Cmax from the same IT dose levels were 120, 418 and 1,103-fold greater than 10 mg/kg PO, and 4, 13 and 34-fold greater than the 100 mg/kg PO dose.

To further characterize the compartmental location of lung-delivered drug, BALF was collected at two time points corresponding with lung tissue sampling (Figure 1). Results were corrected for recovery efficiency and then converted to μg/mL in epithelial lining fluid (ELF) (for comparative purposes only, total lung ELF volume was considered 5 μL). Results indicate ELF nintedanib concentrations following IT administration were greater than total lung tissue yet eliminated at a statistically indistinguishable rate (slop = -4.02 μg/mL/h and -11.03 μg/g/h, respectively; differential p=0.969). Interestingly, while high dose PO administration showed prolonged and increased nintedanib partitioning into lung tissue (slope = +2.74 μg/g/h), an opposite trend was observed in ELF where over the same period nintedanib levels quickly declined (slope −2.37 μg/mL/h; p=0.040).

**Figure 1.**
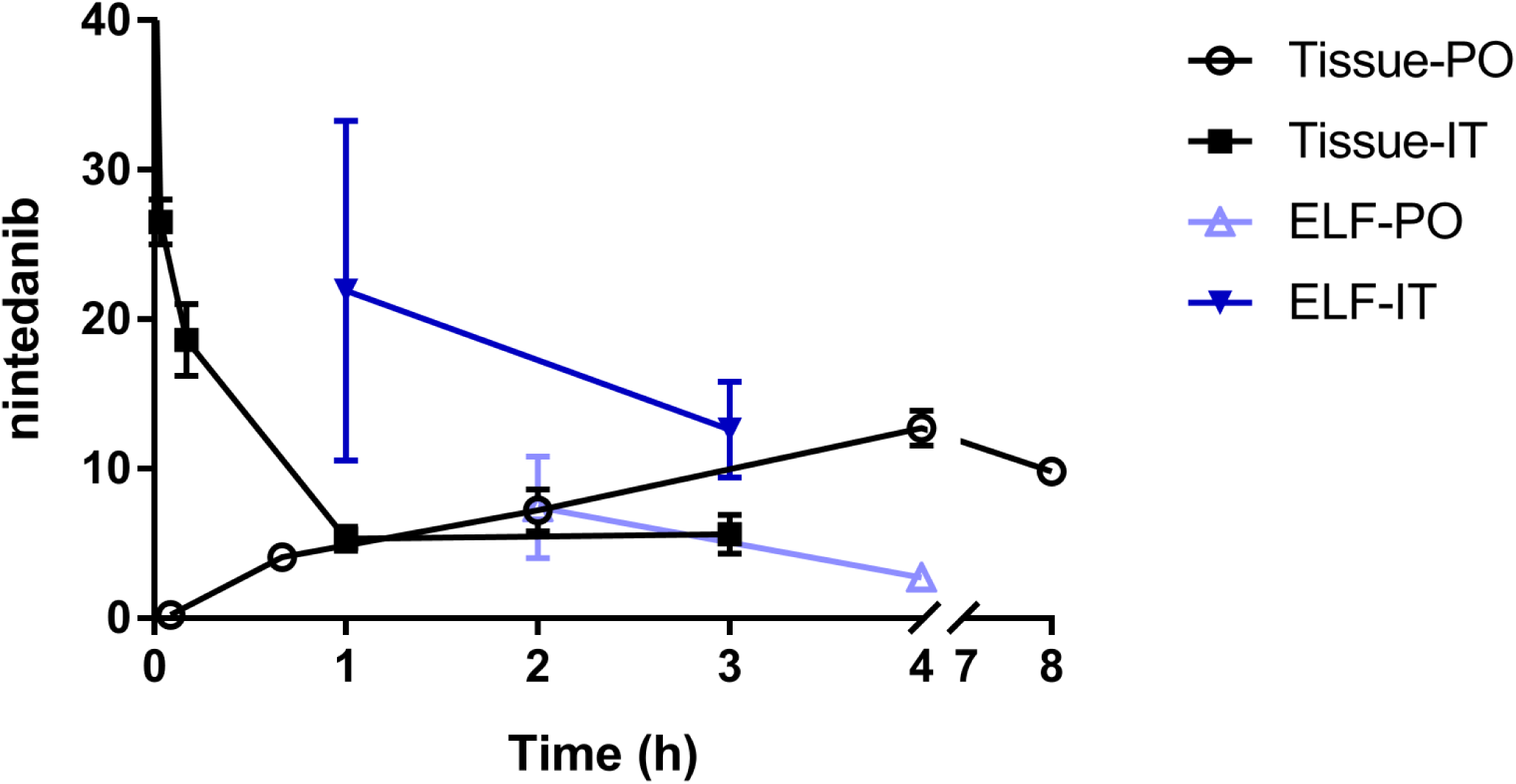
Nintedanib Pharmacokinetics and Lung-Tissue Distribution. Plasma and lung tissue samples were taken 2, 5, 10, 20, 40 min and 1, 2, 4 and 8 hours post dose. For PO administration, nintedanib esylate was dosed in 10 mL/kg at 100 mg/kg (n=3). For Intratracheal (IT) administration, nintedanib salt (1 mg/kg) was dosed in 50 μL at 1 mg/kg (n=9) directly to the lung by nebulizing catheter. BALF was collected at two time points corresponding with lung tissue sampling. BALF was corrected for recovery efficiency and then converted to epithelial lining fluid (ELF) (for these comparative purposes, total lung ELF volume was considered 5 μL). Results represent nintedanib concentration in homogenized lung tissue (μg/g) and ELF (μg/mL).

### 3.2 Short-duration nintedanib exposure inhibits IPF-conditioned matrix effect on normal HLFs

Previously ^[17]^, it was found that exposing normal HLFs (nHLFs) to IPF-conditioned matrix (IPF-CM) increased their aggregate size and consequentially lower aggregate numbers. In addition, IPF-CM exposure resulted in elevated nHLF cell counts. In that work, continuous 24 h nintedanib (100 nM) exposure blocked these pro-fibrotic effects. Here, we initially repeated these published observations and then expanded the dose range to include lower doses (1-10 nM). To explore the nintedanib pharmacokinetic/pharmacodynamic relationship, we compared the effects of short-duration (inhalation mimic) and continuous nintedanib exposure (oral mimic).

To control for interfering wash effects, nHLFs were initially exposed to 1-100 nM nintedanib for 60 min, washed, re-exposed to matching nintedanib (1-100 nM), and incubated for an additional 24 h. As seen previously ^[17]^, results showed nintedanib effectively reduced IPF-CM-induced changes in aggregate size and cell count (Figure 2A and 2C). From these observations, 1 and 10 nM nintedanib were selected for short duration exposure experiments. In these experiments, nHLFs were exposed to 1 and 10 nM nintedanib for 60 min, washed and then incubated without additional nintedanib for 24 h. Results show that short duration, 60 min nintedanib exposure was as effective in reducing IPF-CM induced aggregate size as continuous exposure. Short duration nintedanib exposure (10 nM) also had a similar effect blocking cell count increases, with 1 nM exhibiting a lesser effect (Figure 2B and 2D).

**Figure 2.**
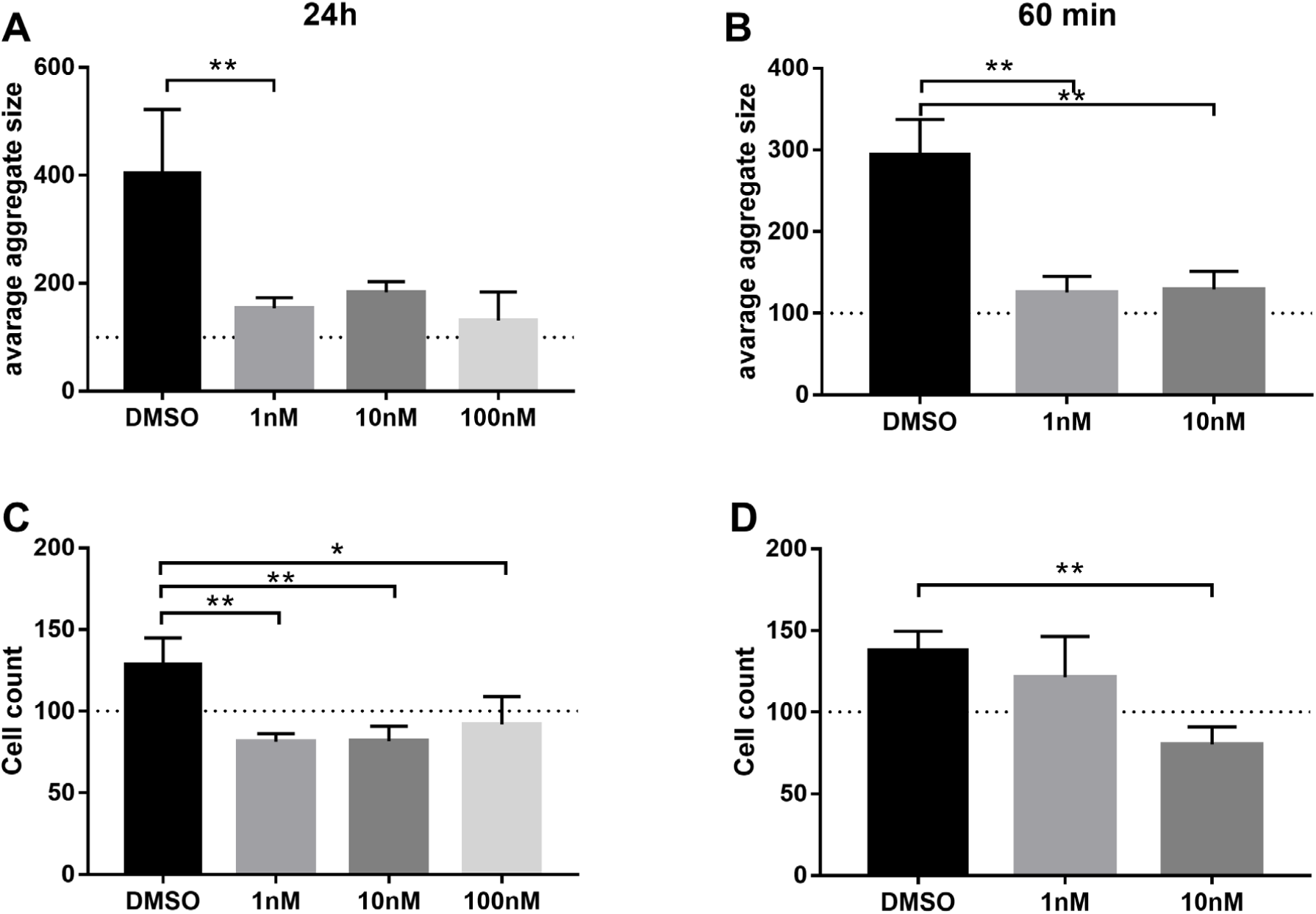
IPF-CMs’ pro-fibrotic activity on normal HLFs is inhibited by nintedanib. Normal human lung fibroblast (nHLFs) were exposed to nintedanib (1-100 nM) for 60 min, washed and further cultured on control or IPF cellular matrix (IPF-CM) with or without nintedanib (1-100 nM) for the entire 24 hr culture incubation (A and C) or only during the initial 60 min (B and D). Following culture, cell distribution was visually evaluated (A and B), and cells were harvested and counted (C and D). Results were normalized to control normal cellular matrix (N-CM). *p<0.05, **p<0.01, n≥4.

IPF-CM exposure was also shown to increase pro-fibrotic signaling, such as Collagen 1a and αSMA ^[17]^. To compare the effects of short duration and continuous nintedanib exposure on these observations, RNA was extracted following the above cultures. Results show that both short-duration (Figure 3B and 3D) and continuous nintedanib (Figure 3A and 3C) exposure inhibited IPF-CM induced COL1A (collagen 1a gene) and ACTA2 (αSMA gene) expression. Interestingly, while COL1A inhibition followed a classical dose response, inhibition of ACTA2 appeared more effective at lower nintedanib concentrations. Similar to changes in aggregate size and cell count, these results show that short duration exposure is as effective as continuous exposure in this human IPF derived *in vitro* model.

**Figure 3.**
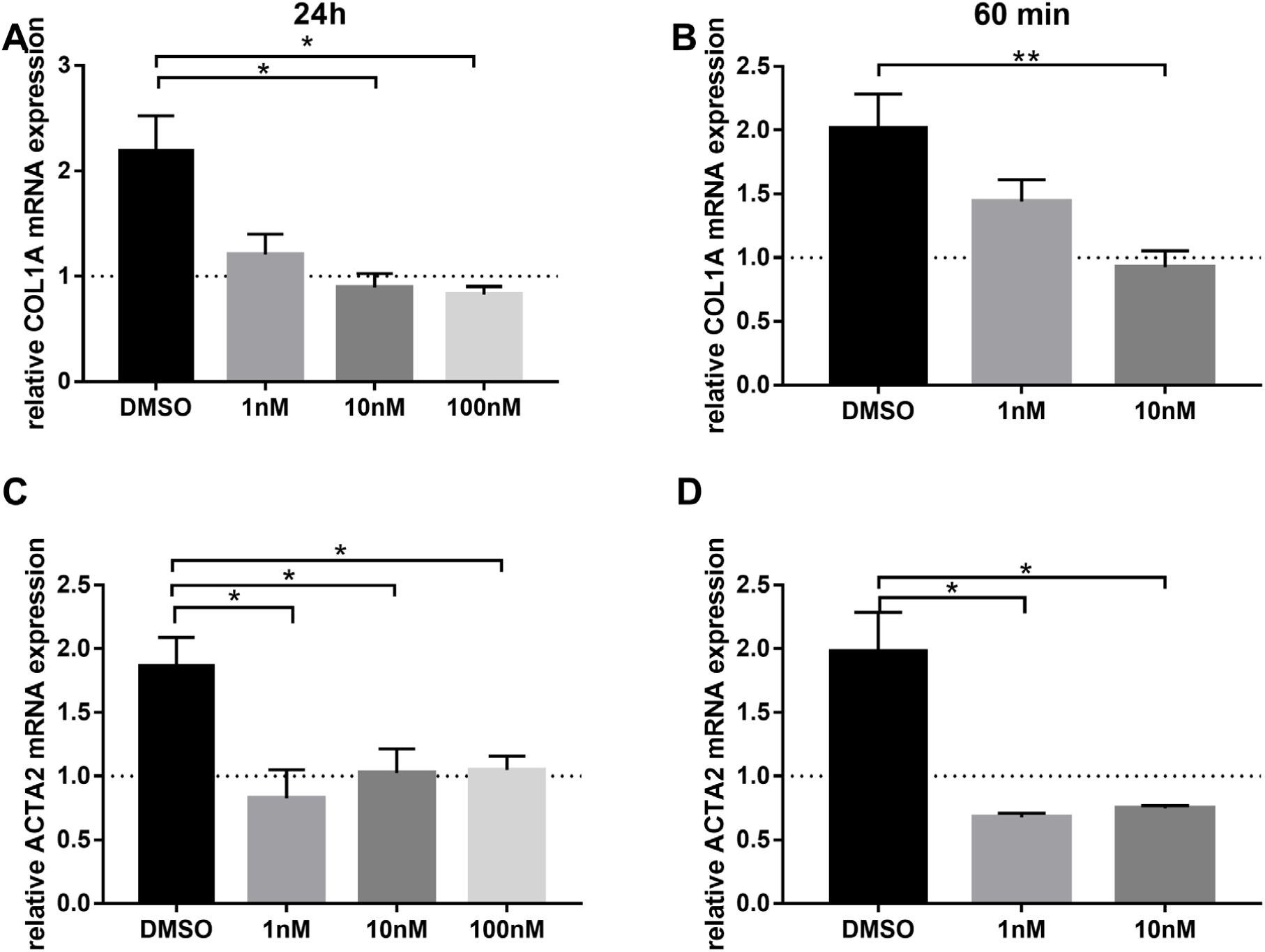
IPF-CMs’ induction of aSMA and Collagen1a in normal HLFs is inhibited by nintedanib. Normal human lung fibroblasts (nHLFs) were exposed to nintedanib (1-100 nM) for 60 min, washed and further cultured on control or IPF cellular matrix (IPF-CM) with or without nintedanib (1-100 nM) for the entire 24 hr culture incubation (A and C) or only during the initial 60 min (B and D). Following culture, RNA was extracted and subjected for qPCR analysis of COL1A (A and B), and αSMA (ACTA2) (C and D). Results were normalized to control normal cellular matrix (N-CM). *p<0.05, **p<0.01, n≥4.

### 3.3 Inhaled nintedanib pharmacokinetics are effective treating silica-induced pulmonary fibrosis in mice

To further characterize the exposure duration required for nintedanib anti-fibrotic activity, the silica-induced pulmonary fibrosis model was performed. Because IT administration is the most efficient method to deliver study medicine to the lung, IT was initially preferred to simulate inhaled, direct-lung delivery. However, unpublished findings showed that bleomycin challenged mice are additionally susceptible to IT administration, making it difficult to separate fibrosis-related events from anesthetic or procedure-associated events (e.g., slow recovery or death from anesthesia and slowed growth rate). Although not bleomycin, to minimize possible non-specific impact of the administration process, we selected IN administration and well-tolerated nintedanib dose levels for this silica study (see Supplement ^[14]^).

As expected, all silica-administered mice lost body weight between days 1-4, then recovered to their initial weight by day 10 (first therapeutic intervention). Body weights remained stable until termination on day 30 (Figure 4A and 4B). No treatment or vehicle administrations altered body weight or disrupted eating habits. Lung function was assessed during termination. While elastance changes between silica-vehicle administered PO or IN mice and sham animals were not significant (Figure 5A), treatment results indicate that both PO and IN nintedanib administration improved lung function, with the latter being dose responsive and the high IN dose achieving significance (p=0.0064; Figure 5A).

**Figure 4.**
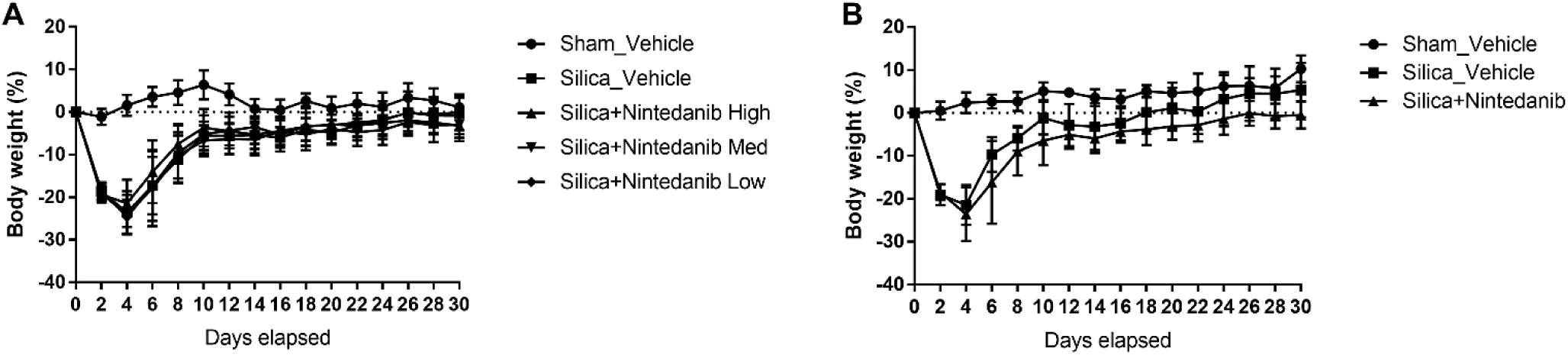
Mouse body weights following silica exposure. Percent body weight following a single silica or PBS IT instillation. Vehicle or nintedanib treatment IN (A) or PO (B) initiated on day 10 and continued through day 29. Body weights collected from time zero through day 30.

**Figure 5:**
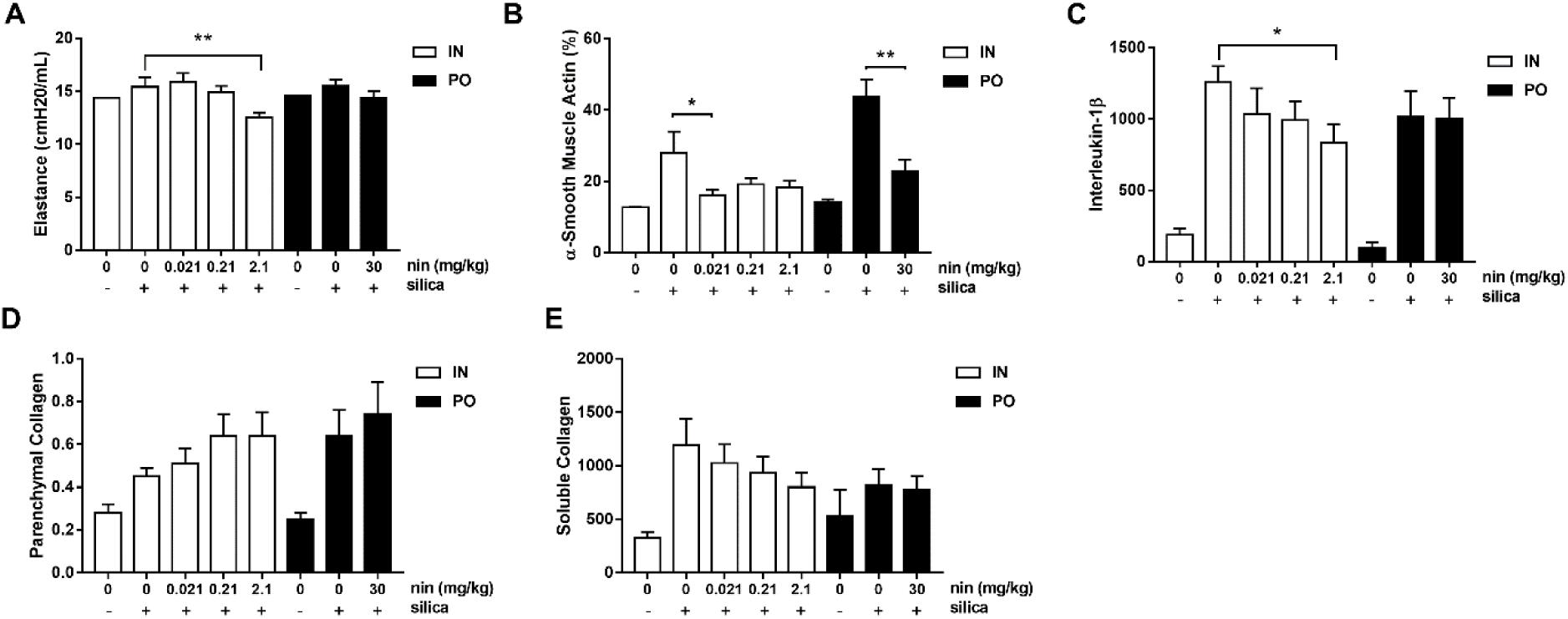
Inhaled nintedanib pharmacokinetics delivers therapeutic anti-fibrotic effects in a mouse Silica model of pulmonary fibrosis. Mice were administered nintedanib by oral gavage (PO, black bars) or inhaled (intratracheal aerosol; IT, white bars) delivery with vehicle (n=10) or nintedanib (n=13). Lung function (elastance) was assessed 30 days post-silica exposure (A). Left lungs were stained to assess and alpha-SMA (B) and parenchymal collagen (D). Right lungs were assessed for interleukin-1b (IL-1b) (C) and soluble collagen (E). *p<0.05, **p<0.01.

#### 3.3.1 Inhaled nintedanib reduces silica-induced pulmonary fibrosis markers in mice

To measure treatment benefit of IN and PO nintedanib in reducing silica-induced fibrotic markers, left lungs were stained for parenchymal collagen and αSMA, while right lungs were analyzed for IL-1β and soluble collagen.

αSMA is a marker of fibroblast differentiation into collagen producing myofibroblasts. Compared to sham animals, silica exposure stimulated a significant increase in lung αSMA expression in IN and PO vehicle-treated mice (Figure 5B). When basal levels were subtracted, QD IN nintedanib treatment exhibited a 78.1, 58.3 and 63.3% benefit for low, mid and high IN dose levels, respectively, with the low IN dose achieving significance (p=0.03; Figure 5B). BID PO nintedanib was also significant (p=0.003; Figure 5B).

Interleukin-1β (IL-1β) is a cytokine important for the initiation and progression of fibrosis. Compared to sham animals, silica exposure stimulated a significant increase in lung IL-1β levels in IN and PO vehicle-treated mice (Figure 5C). When basal levels were subtracted, QD IN treatment exhibited a 22.1, 26.3 and 46.7% dose-responsive benefit for low, mid and high IN dose levels, respectively, with the high IN dose achieving significance (p=0.044; Figure 5C). BID PO nintedanib showed no benefit.

Parenchymal collagen was measured by PSR staining. Results show that IN and PO sham mice exhibited similar PSR staining. Silica-exposed PO vehicle treated mice exhibited 42% more parenchymal collagen than IN vehicle animals (Figure 5D). However, although parenchymal collagen was increased by silica exposure, it was not reversed by either IN or PO nintedanib treatment.

To assess the effect of silica exposure on lung weight and soluble collagen production, right lungs of each mouse were analyzed. Silica exposure significantly (p=0.0021 and p=0.0013, respectively) increased right lung weights in IN-vehicle and PO-vehicle treated mice (0.29 g and 0.26 g, respectively) compared to sham animals (0.18 g and 0.14 g, respectively). Similar to the parenchymal collagen results, neither IN or PO nintedanib treatments significantly reduced elevated lung weights. (0.29, 0.28 and 0.30 g respectively for low, mid and high dose IN animals and 0.28 g for PO).

Compared to sham animals, analysis showed silica exposure significantly increased soluble collagen levels in IN and PO vehicle-treated mice (Figure 5E). QD IN nintedanib showed a dose-responsive, yet insignificant treatment effect. While not achieving significant, benefit analysis showed 20, 30 and 60% dose-responsive improvement across these IN dose groups. BID PO nintedanib showed an insignificant 17% benefit reducing soluble collagen.

Together, these results show infrequent, short duration IN nintedanib lung exposure is as effective inhibiting silica-induced fibrosis markers as more frequent and prolonged PO exposure.

#### 3.3.2 Inhaled nintedanib reduces silica-induced pulmonary inflammatory markers in mice

To further characterize the effect of exposure duration on nintedanib activity, the pulmonary inflammatory index was measured. Results show silica exposure increased cellular density in IN and PO vehicle-treated mice compared to sham animals (Figure 6A). While both QD IN and BID PO active dose groups had a similar resulting index, due to the increased PO vehicle control value, only the PO dose showed benefit (38%).

**Figure 6:**
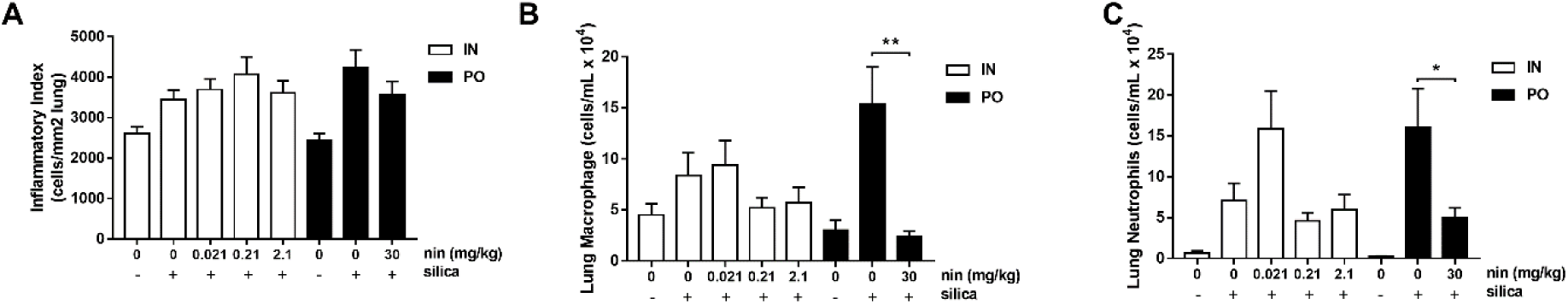
Inhaled nintedanib reduces silica induced inflammation in mouse silica model. Mice were administered nintedanib by oral gavage (PO, black bars) or inhaled (intratracheal aerosol; IT, white bars) delivery with vehicle (n=10) or nintedanib (n=13). The total number of cells per square mm lung tissue section was used as an inflammatory index 30 days post-silica exposure (A). Differential cell counts were performed on the collected BALF for macrophage (B) and neutrophil count (C). *p<0.05, **p<0.01.

Cell differentials showed silica increased BALF macrophage and neutrophil numbers in IN and PO vehicle-treated mice compared to sham animals (Figure 6B-C). Similar to inflammatory index, while both QD IN and BID PO active therapies reduced each cell population to similar numbers, due to increased cell numbers in the PO vehicle control, only PO therapy was determined significant. Silica exposure also increased the number of BALF lymphocytes in IN and PO-vehicle treated mice compared to sham animals. Neither PO or IN nintedanib reduced this effect. BALF eosinophil count was not affected by the silica treatments, nor by the PO or IN nintedanib (data not shown).

## 4. DISCUSSION

Marketed as Ofev^®^, oral nintedanib slows human IPF disease progression. However, only the highest studied clinical dose was shown effective (150 mg BID) ^[11]^ and substantial side effects prevented further dose escalation for possible additional efficacy ^[11, 12]^. Complicating matters, first-pass metabolism and safety-driven dose reduction/stoppage protocols further reduce the delivered lung dose and interrupt required maintenance therapy ^[19]^. To overcome these shortcomings and maximize effect, nintedanib was reformulated as a solution for nebulization and inhaled, direct-lung administration. By inhalation, small inhaled doses largely avoid the gastrointestinal tract and first-pass metabolism, and deliver high local lung concentrations that result in low systemic exposure. With these properties, inhalation is anticipated to circumvent oral-associated side effects and allow local lung dose escalation to test for additional efficacy. To explore this possibility, we used inhalation as a tool to improve our understanding of the key pharmacokinetic parameters required for oral nintedanib activity and better predict inhaled nintedanib effectiveness treating IPF.

Key oral nintedanib pharmacokinetic observations include: 1. Delivered plasma and lung tissue concentrations were disproportionate to dose levels. Despite the 100 mg/kg dose being only 10-fold greater than the 10 mg/kg dose, delivered plasma AUC and Cmax were 55 and 80-fold greater, while lung AUC and Cmax were both ∼34-fold greater; 2. Lung tissue partitioning efficiency decreased with increased dose, suggesting a lung compartment capable of saturation. Specifically, lung AUC and Cmax following the 10 mg/kg dose were 23 and 33-fold higher, respectively than plasma, yet only 13 and 14-fold higher, respectively following the 100 mg/kg dose; and 3. Lung tissue absorption and elimination half-life are longer than plasma. Compared to plasma, the 10 and 100 mg/kg oral dose levels exhibited a 19% and 37% longer lung Tmax and 1.7 and 2.2-fold longer lung terminal half-life, respectively. Taken together, oral nintedanib plasma exposure and lung tissue partitioning were not dose-proportional and delivered lung levels were substantially higher than blood. This data supports previous observations that oral-delivered nintedanib extensively partitions into tissues ^[20]^ and exhibits flip-flop pharmacokinetics ^[21]^, whereby plasma elimination (whether metabolic or tissue partitioning) is faster than absorption (whether continued absorption from the gastrointestinal tract or nintedanib returning to plasma from previously partitioned tissue levels). By performing this analysis at two PO doses, this data provides insight into pharmacokinetic events important for nintedanib activity and indicates that measured plasma levels under-estimate nintedanib systemic exposure following oral administration; an effect also observed for other medicines ^[22]^.

To further characterize these events, direct-lung IT administration was used as a method to separate lung pharmacokinetics from oral-specific events such as gastrointestinal absorption, first-pass metabolism and blood exposure occurring prior to lung partitioning. Analysis showed that lung terminal half-life following both oral and IT administration were equivalent at the lower dose levels yet increased ∼20% at the higher dose levels. Comparatively, while plasma terminal half-life was ∼60 min for both oral doses, plasma terminal half-life following IT administrations were closer to their respective lung tissue terminal half-lives. Interestingly, plasma terminal half-life also increased ∼20% at the higher IT dose level. Comparing these data suggest excess nintedanib associated with higher oral doses partitions to the lung tissue less efficiently than lower dose levels, and regardless of administration route, higher doses were eliminated from the lung more slowly than those delivered from smaller dose levels.

Also noteworthy is that direct-lung administered nintedanib exhibited a plasma terminal half-life more than 2-fold longer than oral. One explanation may be that by-passing the gut fails to activate first-pass metabolic mechanisms, and in turn permits longer nintedanib circulation and higher accumulated blood levels. However, this may in part be explained by lung elimination of the inhaled drug defining plasma half-life (being the primary source of plasma nintedanib). It should be considered that because direct-lung administered nintedanib delivers very high initial lung levels (all IT doses delivered a lung Cmax greater than both tested PO dose levels), the apparent saturation effect observed when increasing the oral dose from 10 mg/kg to 100 mg/kg may be greater following direct lung delivery and in turn slows plasma-absorbed nintedanib from re-partitioning this excess nintedanib back into the inhaled drug-saturated lung. While these events may occur, because nintedanib partitions into many tissues ^[20]^, this alone does not explain the extended plasma half-life, supporting the hypothesis that by-passing metabolic activation may be the greatest contributor.

To identify a lung component capable of saturation at higher dose levels, we measured ELF nintedanib concentrations following IT and PO administration. Results indicate ELF nintedanib concentrations following IT administration were greater than total lung tissue homogenates yet eliminated at a statistically indistinguishable rate (−4.02 μg/mL/h and -11.03 μg/g/h; p=0.969). By comparison, 120 min following a 100 mg/kg PO administered dose, ELF nintedanib concentrations were equivalent to their respective lung tissue homogenates. Surprisingly, while over the next two hours lung tissue nintedanib concentrations continued to increase (slope = +2.74 μg/g/h), ELF concentrations quickly declined (slope = −2.37 μg/mL/h; differential p=0.040). While these results do not exclude ELF as a saturating compartment, they raise the interesting possibility that while oral-administered nintedanib has efficient access to the lung, not all lung-partitioned nintedanib may have access the epithelial airway surface. Because the IPF disease initiates and progress at airway and alveolar epithelial surfaces and through their interaction with underlying fibroblasts ^[3, 4, 23]^, these data suggest that although initially-absorbed oral nintedanib efficiently partitions into the lung as a whole, only a limited, quickly eliminated fraction of the partitioned dose may be available to reach this therapeutically-important compartment. However, despite this apparent capacity and ELF exposure-duration limit, oral nintedanib is effective in slowing IPF progression ^[11, 12]^. One explanation may be that short-duration nintedanib exposure (dose portion efficiently partitioned to ELF) is sufficient for IPF efficacy and the remaining lung-partitioned dose may be relegated to an alternative compartment not available for additional pulmonary anti-fibrotic effect. If true, nintedanib activity may be peak concentration dependent rather than time over AUC, per se.

To explore this further, we followed two approaches to characterize the duration of target exposure required for nintedanib activity. First, we exploited the observation that nintedanib inhibits IPF-CM induced changes in nHLF aggregation and cell count, and ACTA2 and COL1A expression ^[17]^. Using this human IPF *in vitro* system, the duration of nintedanib exposure required to inhibit these pro-fibrotic events was measured. Second, we used the mouse silica-induced pulmonary fibrosis model as a tool to test the exposure-duration requirement *in vivo* (oral vs. inhaled pharmacokinetics).

Important for dissecting lung-exposure requirements, while oral nintedanib delivered a large systemic exposure resulting in a lung Tmax ∼4 h with a broad AUC, inhaled administration delivered an immediate lung Tmax with a more-narrow AUC. Thus, by comparing short duration and continuous exposure *in vitro*, and inhaled and oral activity *in vivo*, we can begin to understand the nintedanib pharmacokinetic/pharmacodynamic relationship as it pertains to the exposure requirements for blocking fibrotic processes.

Aggregate size and single cell number were previously described as markers of fibroblast migration and differentiation ^[24]^. The IPF-CM elevates fibroblast aggregate size and cell count following 24 h exposure. Treating cells with nintedanib, both short duration (inhalation mimic) and continuous exposure (oral mimic) significantly reversed these processes. Compared to several *in vitro* studies conducted with nintedanib at excessively high concentrations ^[25-27]^, the IPF-CM assay demonstrated drug activities in the range of clinical exposure ^[12]^.

For *in vivo* efficacy, the silica-induced pulmonary fibrosis model was employed. All silica-exposed mice grew as expected, with no observed effect from nintedanib or vehicle treatment. Elastance results indicated that both BID oral and QD inhaled nintedanib improved lung function, with the latter being dose responsive and the high inhaled dose achieving significance. Compared to experience in other fibrosis models (e.g., bleomycin ^[28]^ and AdTGFβ ^[29]^), only a small elastance increase was observed in silica treated mice. With this caveat, and the observation that the inhaled-treatment benefit extended elastance improvement beyond sham animals, the clinical relevance of these results should be carefully considered.

During normal healing conditions, fibroblasts differentiate into myofibroblasts with increased αSMA expression and upregulated collagen synthesis. Once the wound is healed, myofibroblasts usually undergo apoptosis ^[30]^. However, when this mechanism becomes dysregulated, chronically activated myofibroblasts can lead to progressive fibrosis ^[31, 32]^. Similar to IPF-CM stimulation of nHLFs, silica exposure induced elevated lung fibroblast αSMA levels and both BID oral and QD inhaled nintedanib reversed this effect. Interestingly, while all inhaled dose levels reduced αSMA, only the inhaled low dose and oral achieved significance. In an unpublished study, these inhaled nintedanib dose levels were separately co-administered with pirfenidone. Similar to the inhaled stand-alone nintedanib result, only the low nintedanib dose achieved significance (p=0.049). These results are also similar to nintedanib inhibition of IPF-CM induced αSMA expression, where lower doses appeared most effective.

As an important cytokine for IPF initiation and progression, IL-1β plays a central role in normal healing and dysregulation of that process ^[33]^. In the fibrotic setting, evidence suggests continuous IL-1β production and subsequent airway/alveolar epithelial cell-to-fibroblast or fibroblast-to-fibroblast signaling could maintain the pro-fibrotic myofibroblast phenotype ^[3]^. *In vivo* silica study results showed that QD inhaled nintedanib was dose responsive and significant in reducing IL-1β levels. While others have shown this oral dose level effective in this model ^[10]^, PO administration in this study did not significantly reduce IL-1β.

Although inhaled nintedanib was dose responsive inhibiting soluble collagen, neither changes in parenchymal collagen or increased lung weights were reversed by oral or inhaled nintedanib. Due to the positive impact on key fibrotic markers (αSMA and IL-1β, and soluble collagen) these results were surprising. As an explanation, it is possible that treatment duration or the nintedanib mechanism itself may be insufficient to reverse established fibrosis ^[34]^; a hypothesis supported by clinical observations that Ofev treatment only slows annual FVC decline, rather than reversing fibrosis ^[12, 35]^.

The effect of nintedanib on inflammation was also measured. Results show that both BID oral and QD inhaled nintedanib reduced silica-induced macrophage and neutrophil numbers, with the former achieving significance. These findings support a recent study showing that BALF neutrophils are reduced following nintedanib treatment in this model ^[6]^. Interestingly, inhaled control groups exhibited a smaller silica-induced increase in macrophage and neutrophil numbers compared to oral. Because the pro-fibrotic signals were similar across all silica-exposed animals, these results suggest a possible vehicle or anesthetic anti-inflammatory effect.

In summary, this report provides data supporting oral flip-flop pharmacokinetics showing measured plasma levels underestimate nintedanib lung tissue exposure. These data also suggest only a small fraction of the oral dose may be delivered to the pulmonary epithelial surface and despite continued tissue partitioning, may only exist in this compartment for a short duration. Because this compartment is important for IPF initiation and progression, these observations suggest that only relatively short duration nintedanib exposure may be required for clinical efficacy. In support of this possibility, both human IPF *in vitro* and *in vivo* animal models demonstrated that only short duration exposure (inhaled pharmacokinetics delivered QD) were required for nintedanib activity.

As shown here and supported elsewhere ^[8, 13]^, pharmacokinetic elements important for nintedanib activity can be delivered by infrequent, small inhaled doses to achieve oral equivalent-to-superior activity in the lung. While additional studies are required to confirm the differential impact of these key pharmacokinetic parameters in treating IPF, the possible patient benefit generated by inhaled nintedanib (i.e. reduced side effects and possibility to increase functional lung-delivered nintedanib for improved efficacy) should motivate further research.

## Acknowledgements

This work was funded by Avalyn Pharma, Inc., Seattle, Washington 98101, United States

## CRediT Authorship Contribution Statement

MS Conceptualization, Design, Formal analysis, Writing, review and editing; GES Investigation, Formal analysis, Report generation, Writing; SP Investigation, Report generation; SB Data review; SA, OM, SR, AH, MV, BBW Investigation, Report generation; DS, KA, ABM and MK Supervision, Analysis, Report generation. All authors read and approved final version of manuscript.

