## Supplement for "Inhalation: A means to explore and optimize nintedanib’s pharmacokinetic/pharmacodynamic relationship"

a. Meir Medical Center, Pulmonary Department, Kfar Saba, 4428164, Israel
b. Tel Aviv University Sackler Faculty of Medicine, Tel Aviv 6997801, Israel
c. Avalyn Pharma, 701 Pike Street, Suite 1500, Seattle, WA 98101, United States
d. McMaster University, Hamilton, ON, L8S 4L8, Canada
e. Firestone Institute for Respiratory Health, Hamilton, ON, L8N 4A6, Canada

**Corresponding author:**

Mark W. Surber

### METHODS

**Table S-1. Study dose solution compositions**

| Route | Dose Level | Dose Solution Composition |
|---|---|---|
| Oral (PO) | Vehicle | 1% methylcellulose |
| | 10 mg/kg nintedanib | 1.0 mg/mL nintedanib<br>1% methylcellulose |
| | 30 mg/kg nintedanib | 3.0 mg/mL nintedanib |
| | 100 mg/kg nintedanib | 10 mg/mL nintedanib<br>1% methylcellulose |
| Inhaled (IT) | 1 mg/kg | 0.4 mg/mL nintedanib<br>2.0% propylene glycol |
| | 2.5 mg/kg | 1.0 mg/mL nintedanib<br>2.0% propylene glycol |
| | 10 mg/kg | 4.0 mg/mL nintedanib<br>2.0% propylene glycol |
| Inhaled (IN) | Vehicle | 1.5% propylene glycol<br>0.4% NaCl |
| | 0.021 mg/kg nintedanib | 0.012 mg/mL nintedanib<br>1.5% propylene glycol<br>0.4% NaCl |
| | 0.21 mg/kg nintedanib | 0.12 mg/mL nintedanib<br>1.5% propylene glycol<br>0.4% NaCl |
| | 1.5 mg/kg nintedanib | 0.9 mg/mL nintedanib<br>1.5% propylene glycol<br>0.4% NaCl |
| | 2.1 mg/kg nintedanib | 1.2 mg/mL nintedanib<br>1.5% propylene glycol<br>0.4% NaCl |
| | 3.0 mg/kg nintedanib | 1.7 mg/mL nintedanib<br>1.5% propylene glycol<br>0.4% NaCl |

**Intranasal/intratracheal pharmacokinetic bridging btudy**

For IN pharmacokinetic analysis, groups of 16 female C57BL/6 mice were placed into four groups of four mice and under light isoflurane anaesthesia, dosed 3.0 mg/kg nintedanib in 50 uL water with 1.5% propylene glycol and 0.4% NaCl, evenly divided between the two nostrils. Because IN administered aerosol comes in contact with the upper airway, permeant anion was required for IN dosing solution tolerability. Formulated solutions were administered within two hours of compounding. Plasma (in $K_3EDTA$) and lungs were harvested at 2, 10, 20 and 120 min post administration (n of 3 per time point per group). All non-terminal and 2-minute terminal blood samples were taken from fascial bleeds. All other terminal bleeds were taken from the inferior vena cava.

**Bioanalytical analysis**

For nintedanib quantitation in plasma, nintendanib-$d_3$ internal standard (I.S.; Toronto Research Chemicals, New York, Canada) was added to plasma samples and subjected to a liquid-liquid extraction procedure. For lung analysis, samples were first homogenized using Lysing Matrix D (MP Biomedical, Irvine, CA). Homogenized lung preparations were extracted and analysed as described for plasma. Pharmacokinetic parameters were determined using the linear trapezoidal method. Briefly, the organic portion containing nintedanib and I.S. were transferred to a clean tube and evaporated under nitrogen. The extract was reconstituted and diluted in methanol:water (50:50, v/v). An aliquot was analyzed by reversed-phase HPLC (Waters, Milford, MA) using a Allure Biphenyl column (Restek Corp, Bellefonte, PA), maintained at 35°C. The mobile phase was nebulized using heated nitrogen in a Z-spray source/interface set to electrospray positive ionization mode. The ionized compounds are detected using MS/MS (Quattro Premier Mass Spectrophotometer; Waters, Milford, MA).

**Tolerability study**

On day one, while under light isoflurane anesthesia, pulmonary fibrosis was induced in eight 20-22 g female C57BL/6 mice by a single intratracheal intubation of silica (2.5 mg/kg) (U.S. Silica, Katy, Texas). On day 10, mice were placed into two groups of four and administered either 3.0 mg/kg or 1.5 mg/kg nintedanib in 50 µL formulation (approximately 25 µL to each nostril). Mice were then placed into a separate cage and monitored for at least two hours post dose. All mice received a second IN nintedanib dose 24 hours later and again observed for 2 hours, followed by a 7-day observation period.

**Lung function assessment**

Flexivent determined elastance was used as a measure of lung stiffness/fibrosis by inflating the lungs and assessing changes in flow and pressure as described previously (S-1, S-2). Briefly, 30 days following silica instillation, mice were sedated with an intraperitoneal injection (IP) of xylazine hydrochloride (10 mg/kg) followed by IP injection of ketamine (150 mg/kg). A tracheotomy was performed, and a 19-gauge blunted needle was inserted into the trachea; the mice were attached to a rodent mechanical ventilator (Flexivent, SCIREQ). In between forced oscillation waveforms maneuvers, mice were ventilated with 10 ml/kg air at a rate of 150 breaths per minute. Changes in flow, volume and pressure within the airways was recorded and the raw data fit to a single compartment model to assess quasi-static elastance using the P-V Loop *Salazar-Knowles equation*.

**BALF collection for cytospins**

After ketamine/xylazine anesthesia, the thoracic cavity was opened and mice were euthanized via exsanguination. The lungs from each mouse were excised and sequentially lavaged with 600 µL PBS followed by 400 µL PBS using a 1 mL syringe and cannula. For each wash, lungs were briefly massaged to harvest lung infiltrate cells. Each PBS lavage was extracted using the same 1 mL syringe. Recovered bronchoalveolar lavage fluid (BALF) from each mouse was pooled (~400-500 µL) and placed in a 1.5 mL Eppendorf tube. Total cell count (in 20 µL lavage fluid per sample) was performed and trypan blue stain 0.4% (Invitrogen, cat#

T10282) added to test cellular viability. BALF samples were centrifuged at 14,000 rpm for 5 min. Supernatant was stored at –80°C and the pellet suspended in cold PBS (600 µL-1000 µL). 120 µL of each sample was used to prepare smears using cytocentrifugation (Shandon, Pittsburgh, PA, USA; 1000 rpm for 3 min). Developed cytospin slides were sent to the McMaster University histology core facility for Wright–Giemsa staining according to the manufacturer's protocol. Cell differentials were determined from approximately 200 random counts using a hemocytometer. Cells were classified as neutrophils, macrophages, eosinophils or lymphocytes.

**Lung processing for histopathology and sircol assay**

Lung histopathology and Sircol assay were performed as described previously (S-2). Following BALF collection, a surgical suture was used to tie off and separate the left lung lobe from the four right lobes. The four right lung lobes were snap frozen in liquid nitrogen and stored at -80ºC. The frozen right lung lobes were weighed and placed in a metal container. A piston was inserted and the top hammered to crush frozen lung lobes into a fine powder. Powdered lungs were removed, weighed and transferred into tubes for Sircol collagen assay (see below). The left lobe of each lung was inflated to 30 cm$H_2O$ for 3–5 min in 10% formalin solution for paraffin block preparation.

**Sircol soluble collagen assay**

Crushed lung powder was homogenized in 1 mL RIPA buffer. Homogenates were centrifuged at 440g for 8 min. Soluble collagen was determined in RIPA-homogenized lung tissues, free from cell debris and insoluble ECM fragments (Sircol™ soluble collagen assay; Biocolor Ltd, Carrickfergus, UK). Results were expressed as µg soluble collagen per mL solution.

**Interleukin-1 beta ELISA**

Interleukin-1 beta (IL-1β) was measured as described previously (S-2). Briefly, IL-1β concentrations were measured in the supernatant from right lung homogenates using commercially available DY401 ELISA kits (R&D Systems, Minneapolis, MN, USA).

**HALO™ analysis**

Immunohistochemical (IHC) stained microscope slides were digitalized using an Olympus VS120-L100-W slide scanner at a 20× magnification and quantified using HALO™ Image Analysis Software through cytonuclear and area quantification modules (Indica Labs, Albuquerque, New Mexico). Fixed lung tissues were processed, embedded in paraffin and stained with H&E, IHC for α-SMA, and with picosirius red (PSR) for parenchymal collagen. Total lung tissue area ($mm^2$) was measured for each specimen. All cells in the tissue area were quantified (total cell count) and any cells containing the brown-stained αSMA were considered positive. PSR-stained lungs were imaged using polarized light (POL-MONO), producing a high-contrast black and white image with white corresponding to collagen fiber. HALO was used to quantify lung parenchymal collagen content (area; $mm^2$). Collagen-rich airways were omitted from the analysis by cross-reference with its associated H&E scan.

### RESULTS

#### Intranasal/intratracheal pharmacokinetic bridging study

Prior to study start, an IN pharmacokinetic bridging study was performed and compared to IT results. It was determined that IN lung delivery was roughly half as efficient at IT administration, exhibiting a ~10-fold longer alpha-phase lung elimination half-life, 2-fold longer lung terminal half-life, and IN plasma levels exhibited similar blood AUC, with roughly 2-fold greater Cmax. These differing results may be explained by the administration route. The IN dose initially passes through the highly vascular upper airway, resulting in some dose loss to the blood and lower lung Cmax than IT, where the entire dose is first delivered to the lung and the majority of resulting blood levels are derived from pulmonary elimination. Despite these differences, IN administration maintained the desired inhaled pharmacokinetic characteristics, wherein an immediate lung Cmax and short duration lung exposure was achieved (Epstein-Shochet et al., 2020 Manuscript, Table 2).

#### Silica model of pulmonary fibrosis – tolerability and dose selection

Previous mouse studies indicated multi-dose IT administration is not well-tolerated (data not shown). To optimize animal health during the multi-dose silica-induced fibrosis study, we explored the tolerability of IN inhaled nintedanib in silica-challenged mice. Results showed no signs of respiratory distress, dyspnea or CNS effects, indicating that IN delivered nintedanib dose levels, drug concentration and formulation were well-tolerated. Due to extra-nasal bubbling of the administered 50 μL dose, the IN volume used in the main silica therapeutic study was reduced to 35 uL (reducing the high dose from 3.0 mg/kg to 2.1 mg/kg). Extrapolating these and IN pharmacokinetic data, three silica fibrosis model dose levels were selected to broadly span the 30 mg/kg oral delivered lung Cmax and AUC. These results are shown in Table S-2 and Figure S-1.

**Table S-2. Silica Study Dose Levels and Comparative IN/PO Pharmacokinetics**

| | | Lung Cmax | | Lung AUC | |
|---|---|---|---|---|---|
| Route | Dose (mg/kg) | (µg/g) | Fold vs. PO | (mg·hr/kg) | Fold vs. PO |
| PO | 30 | 1.2 | – | 7.4 | – |
| IN[a] | 2.1 | 29.3 | 24.4 | 39.8 | 5.4 |
| IN[b] | 0.21 | 2.93 | 2.44 | 3.98 | 0.54 |
| IN[c] | 0.021 | 0.293 | 0.244 | 0.398 | 0.054 |

a. PO-superior lung Cmax and AUC; b. PO-equivalent lung Cmax and AUC; c. PO-inferior lung Cmax and AUC

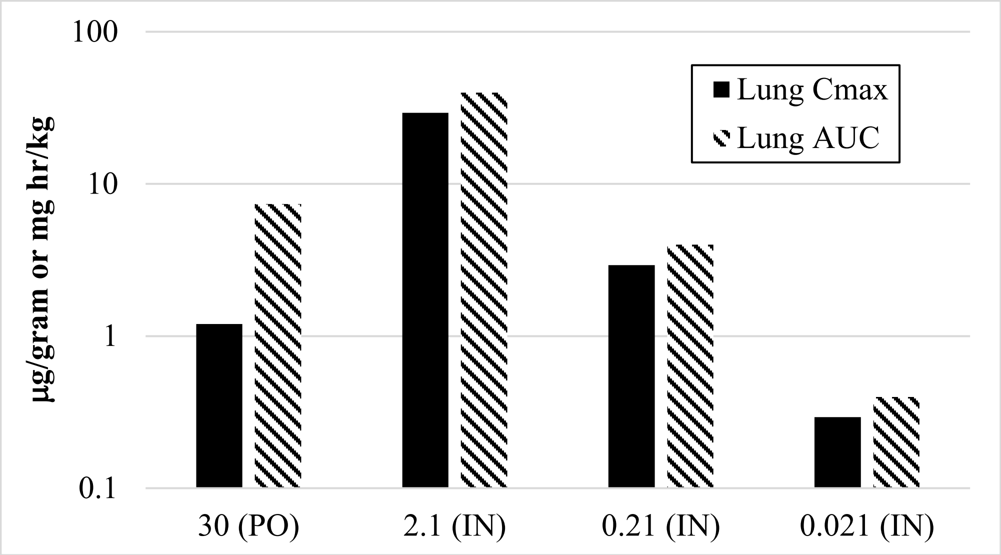


**Figure S-1. Silica Study Dose Levels and IN/PO Comparative Pharmacokinetics**

### REFERENCES

S-1 Ayaub, E. A., Dubey, A., Imani, J., Botelho, F., Kolb, M. R., Richards, C. D., & Ask, K. Overexpression of OSM and IL-6 impacts the polarization of pro-fibrotic macrophages and the development of bleomycin-induced lung fibrosis. Scientific Reports 2017; 7, 13281.

S-2 Ayaub, E. A. Kolb P. S., Mohammed-Ali Z., Tat V., et al. GRP78 and CHOP modulate macrophage apoptosis and the development of bleomycin-induced pulmonary fibrosis. J Pathol 2016; 239, 411–425.
